# Investigating the effects of repetitive paired-pulse transcranial magnetic stimulation on visuomotor training using TMS-EEG

**DOI:** 10.1101/2024.02.21.581468

**Authors:** Ryoki Sasaki, Brodie J. Hand, Wei-Yeh Liao, John G. Semmler, George M. Opie

## Abstract

**Objectives:** I-wave periodicity repetitive paired-pulse transcranial magnetic stimulation (iTMS) can modify acquisition of a novel motor skill, but the associated neurophysiological effects remain unclear. The current study therefore used combined TMS-electroencephalography (TMS-EEG) to investigate the neurophysiological effects of iTMS on subsequent visuomotor training (VT).

**Methods:** Sixteen young adults (26.1 ± 5.1 years) participated in three sessions including real iTMS and VT (iTMS + VT), control iTMS and VT (iTMS*_sham_* + VT), or iTMS alone. Motor-evoked potentials (MEPs) and TMS-evoked potentials (TEPs) were measured before and after iTMS, and again after VT, to assess neuroplastic changes.

**Results:** Irrespective of the intervention, MEP amplitude was not changed after iTMS or VT (*P* = 0.211). Motor skill was improved compared with baseline (*P* < 0.001), but no differences were found between stimulus conditions. In contrast, the P30 peak was altered by VT when preceded by sham iTMS (*P* < 0.05), but this effect was not apparent when VT was preceded by iTMS or following iTMS alone (all *P* > 0.15).

**Conclusion:** In contrast to expectations, iTMS was unable to modulate MEP amplitude or influence motor learning. Despite this, changes in P30 amplitude suggested that motor learning was associated with altered cortical reactivity. Furthermore, this effect was abolished by priming with iTMS, suggesting an influence of priming that failed to impact learning.

**Authorship statements:** Conceptualization: JGS; Data curation: RS, BJH, and WL; Formal analysis: RS; Funding acquisition: RS; Investigation: RS, BJH, and WL; Methodology: RS, GMO, BJH and JGS; Project administration: GMO and JGS; Supervision: GMO and JGS; Roles/Writing - original draft: RS and GMO; Writing - review & editing: BJH, WL, and JGS.

## Introduction

Learning new motor skills is an essential aspect of daily life that is associated with neuroplastic changes in the brain. These changes are characterized by the modulation of existing neural communication and the formation of new connections (for review, see Dayan & Cohen, 2011). This role of neuroplasticity in mediating motor learning means that factors influencing plasticity induction also have the potential to influence the extent of learning. Given the clear benefits of such capabilities in both healthy and pathological populations, an extensive literature aiming to modulate learning by manipulating plasticity has developed (Jung & Ziemann, 2009; Fujiyama *et al*., 2017; Sasaki *et al*., 2018; Opie *et al*., 2020). A popular approach within this literature has been to leverage the concept of metaplasticity, wherein the sign and magnitude of a neuroplastic change is determined by previous activity within the targeted synapses (for review, see Ziemann & Siebner, 2008). Within this construct, an intervention able to produce a directed change in brain activity is applied before a period of training to ‘prime’ neuroplastic changes associated with training (Jung & Ziemann, 2009; Fujiyama *et al*., 2017; Sasaki *et al*., 2018; Opie *et al*., 2020).

The utility of this priming approach has been facilitated in humans by the application of different forms of non-invasive brain stimulation (NIBS). These techniques can induce short-term neuroplastic changes in the brain (Nitsche & Paulus, 2000; Stefan *et al*., 2002; Huang & Rothwell, 2004; Peinemann *et al*., 2004) and have been shown to influence skill acquisition in a metaplastic way (Jung & Ziemann, 2009; Jelić *et al*., 2015; Fujiyama *et al*., 2017; Sasaki *et al*., 2018; Opie *et al*., 2020). Much of the literature investigating the influence of priming NIBS on motor learning has applied more conventional stimulation (e.g., theta burst stimulation [TBS], paired-associative stimulation [PAS], transcranial direct current stimulation [tDCS]). However, we recently demonstrated that I-wave periodicity repetitive paired-pulse TMS (iTMS) – an intervention that targets activity of local intracortical circuits in primary motor cortex (M1) – is also able to facilitate acquisition of a novel motor skill (Hand *et al*., 2023). While this demonstrates the utility of iTMS as a priming tool, this study also found inconsistencies between the neurophysiological and functional response to priming. Consequently, the mechanisms that underpin the functional effects of priming with iTMS remain unclear, which limit application of this approach.

Within the current study, we sought to address this limitation by using TMS in conjunction with electroencephalography (TMS-EEG). Recent work from our group suggests that the TMS-evoked EEG potential (TEP) can reveal central effects of iTMS which are not indexed by motor evoked potentials (MEPs)(Sasaki *et al*., 2023). We therefore reasoned that the TEP may be able to provide some additional neurophysiological insight to how iTMS influences motor learning. Consequently, TEPs were recorded before and after practice of a novel visuomotor adaptation task, either in isolation or following application of real or sham iTMS.

## Methods

### Participants

A total of 16 healthy young adults (7 men and 9 women; mean age ± SD = 26.1 ± 5.1 years; age range = 19–35 years) were recruited from the University and wider community to participate in this study. All participants were right-handed, free of neurological and psychiatric disorders, were not taking any drugs that influence the central nervous system and had normal or corrected-to-normal vision. Contraindications to TMS were assessed using the TMS adult safety screen (Rossi *et al*., 2009). A nominal payment of $15 per hour was offered to compensate for time and cost of participation. Written informed consent was provided prior to inclusion and the study was conducted in accordance with the *Declaration of Helsinki.* All experimental procedures were approved by the University of Adelaide Human Research Ethics Committee (approval number: H-026-2008).

### Experimental Arrangement

All participants attended three experimental sessions that were each approximately 3.5 hours long, held at the same time of day and separated by at least one week (Figure 1). Each session involved recording MEPs and TEPs before (Pre) and immediately after iTMS (Post iTMS), as well as after visuomotor training (VT)(Post Train). Sessions included real iTMS and VT (iTMS+VT), sham iTMS and VT (iTMS_sham_+VT) and iTMS only (iTMS), with the order of sessions randomized within a participant. For each session, participants sat in a comfortable chair with their right hand pronated on a table and were instructed to keep their eyes open and remain relaxed. Surface electromyography (EMG) was recorded from the right first dorsal interosseous (FDI) muscle via disposable Ag/AgCl electrodes in a belly−tendon montage, with an additional Ag/AgCl electrode placed over the right ulnar styloid as an earth electrode. EMG data were sampled at 2000 Hz using a CED1401 interface (Cambridge Electronic Design, Cambridge, UK), amplified (1000×) and band-pass filtered (20–1000 Hz) by a CED1902 signal conditioner (Cambridge Electronic Design, Cambridge, UK). Line noise was removed using a Humbug mains eliminator (Quest Scientific, North Vancouver, Canada) and recordings were stored on a personal computer for off-line analysis.

**Figure 1.**
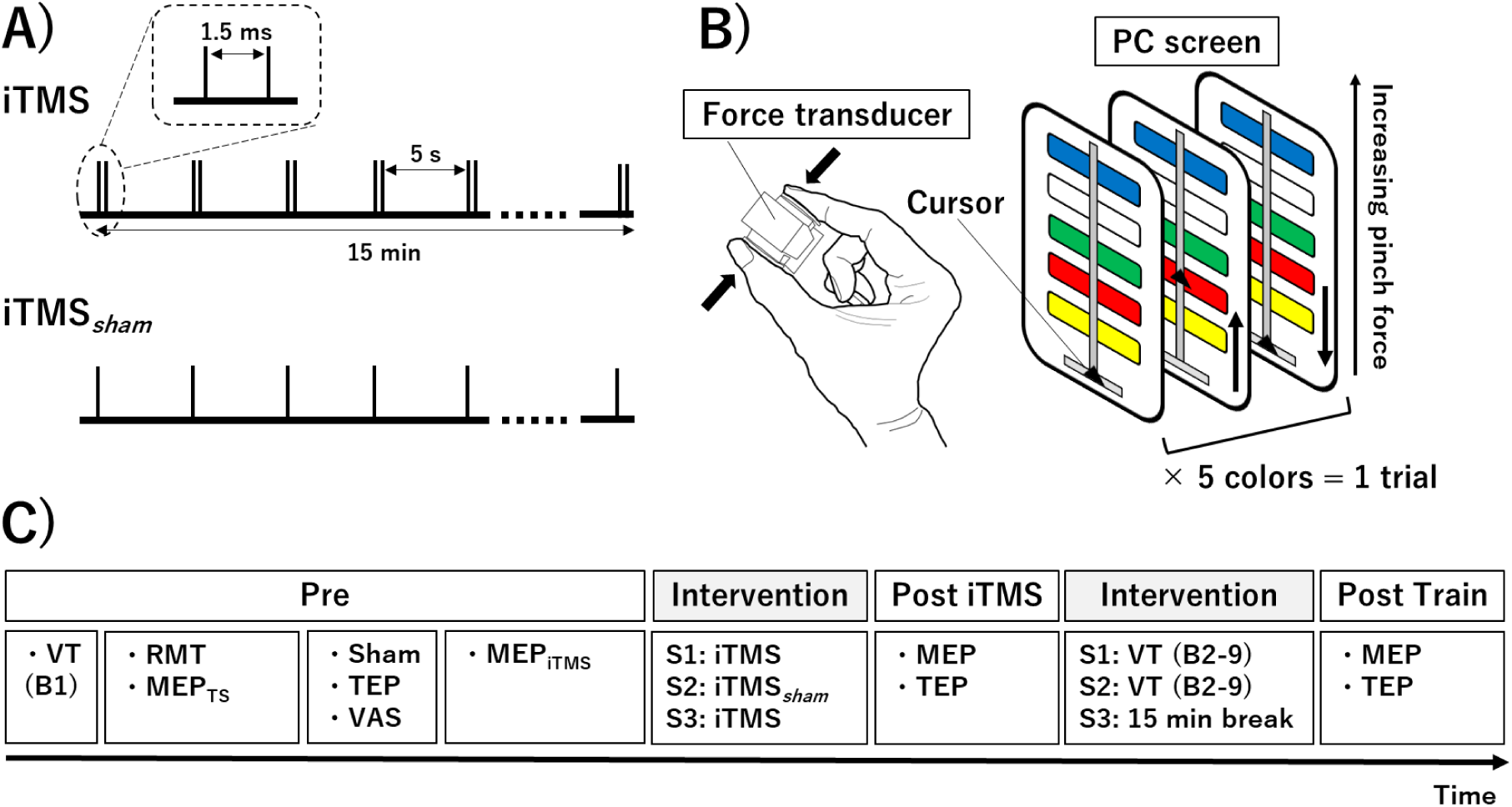
Intervention settings and experimental protocol. (A) iTMS intervention parameters. (B) Visuomotor training setup and requirements. (C) Experimental protocol. Three experimental sessions were performed involving different combinations of iTMS (S1: iTMS; S2: iTMS_sham_; S3: iTMS) and VT (S1: VT; S2: VT; S3: 15 minutes break). Cortical excitability indexed with both MEPs and TEPs (sham and real TMS) was recorded before iTMS (Pre), immediately after iTMS (Post iTMS) and immediately after VT (Post Train). Abbreviations; B, block; iTMS_sham_, control I-wave periodicity repetitive transcranial magnetic stimulation; iTMS, I-wave periodicity repetitive transcranial magnetic stimulation; MEP, motor-evoked potential; MEP_iTMS_, MEP amplitude producing a response of ∼0.5–1 mV by iTMS; MEP_TS_, MEP amplitude producing a response of ∼0.5–1 mV by single pulse TMS; S, session; TEP, transcranial magnetic stimulation-evoked potential; TMS, transcranial magnetic stimulation; VAS, visual analog scale; VT, visuomotor task.

#### TMS

Monophasic TMS pulses were delivered to the hand area of the left M1 using a figure-of-eight branding iron coil connected to two Magstim 200^2^ stimulators via a Bistim unit (Magstim, Dyfed, UK). The coil was held tangentially to the scalp at an angle of approximately 45° to the sagittal plane, at the location producing the largest stable response in the resting right FDI muscle with a posterior–anterior coil orientation. This position was co-registered to the MNI-ICBM152 brain template (Fonov *et al*., 2011) using a Brainsight neuronavigation system (Rogue Research Inc, Montreal, Canada). TMS was applied at a rate of 0.25 Hz for MEP and TEP measures with a 10% jitter between trials. Resting motor threshold (RMT) was defined as the minimum intensity needed to evoke MEPs ≥ 50 µV in 5 of 10 consecutive trials during relaxation of the right FDI muscle (Rossini *et al*., 2015). TMS intensity was expressed as a percentage of maximum stimulator output (%MSO). The test stimulus (TS) for MEP measures was set at the intensity required to produce an MEP of ∼0.5–1 mV (MEP_TS_) when averaged over 15 trials.

#### iTMS

iTMS involved 180 pairs of stimuli applied every 5 s, resulting in a total intervention time of 15 minutes (Opie *et al*., 2018; Opie *et al*., 2021). The intensity was the same for both stimuli (Sasaki *et al*., 2022), and was adjusted so that paired stimulation produced a response amplitude of ∼0.5–1 mV (MEP_iTMS_) assessed over 15 trials before the intervention. An interstimulus interval of 1.5 ms (corresponding to I-wave periodicity) was used. In addition, a sham intervention not expected to modulate cortical excitability (single-pulse TMS for 15 min; iTMS*_Sham_*) was applied in a separate session. To avoid coil heating during the intervention, ice packs were always used to cool the coil prior to and during iTMS application. This ensured that the same coil could be used for all TMS measures.

#### EEG

EEG data was recorded using a WaveGuard EEG cap (ANT Neuro, Hengelo, The Netherlands), with 62 sintered Ag/AgCl electrodes in standard 10-10 positions, connected to an eego mylab amplifier (ANT Neuro, Hengelo, The Netherlands). CPz was used as the reference electrode for all recordings. Signals were filtered online (DC–0.26 × sampling frequency), digitized at 8 kHz, and stored on a personal computer for offline analysis. The impedance of all electrodes was constantly kept <10 kΩ through the experiment.

TEPs were recorded in a single block of stimulation that involved 100 pulses set at an intensity of 100% RMT. In addition, a single block of realistic sham stimulation was also recorded, which was designed to quantify the somatosensory- and auditory-evoked potentials that can confound the direct brain response to TMS (Conde *et al*., 2019). This was achieved by applying an electrical stimulus (ES) to the scalp that was timed to coincide with the application of TMS. To do this, a bar electrode was attached to the face of the TMS coil via a plastic clip (∼3 cm length) and held against the EEG cap over the M1 hotspot. This ensured that the TMS coil was adequately separated from the head, while still allowing coil vibration to contribute to somatosensory input. Intensity of ES was set at 3 × perceptual threshold and stimuli were 0.2-ms square-wave constant-current pulses (DS7AH, Digitimer, UK). Sham stimulation involved application of 100 coincident ES and TMS pulses, with TMS set at 100% RMT. During all EEG recordings, participants listened to white noise played through in-canal earphones, with ear defenders (Peltor Optime, 3M; 34db reduction) to minimize the influence of auditory-evoked potentials. The volume of auditory masking was individually adjusted to minimize audition of the TMS click (Biabani *et al*., 2019; Rocchi *et al*., 2020). The perception of real TMS and control conditions was evaluated after baseline EEG recordings. Participants were instructed to fill out a set of visual analog scales (VAS) rating (from 0 to 10): (1) intensity of auditory sensation; (2) intensity of scalp sensation; (3) area of scalp sensation; (4) intensity of pain or discomfort (Gordon *et al*., 2021).

### Visuomotor Training

A sequential visual isometric pinch task (SVIPT) was used to assess motor skill acquisition (Opie *et al*., 2020; Hand *et al*., 2021; Hand *et al*., 2023). Before the task, maximum voluntary contraction (MVC) force was assessed by a force transducer. Participants grasped the transducer between the right index finger and thumb for 3–5 s (repeated three times). The highest force value was set as MVC. During the task, the position of a digital cursor was manipulated by a participant using the pinch grip, with the aim of a single trial being to accurately move the cursor between 5 color targets in a specific order (consistent within each session) while returning to baseline (0% MVC) between each color. The coloured targets disappeared at the end of each trial and reappeared for the start of the next trial. To increase task difficulty and reduce carry-over of learning between sessions, a non-linear transform was used to relate force application to cursor movement. Specifically, logarithmic, exponential and sigmoidal transforms were used for the iTMS+VT, iTMS*_Sham_*+VT and iTMS sessions, respectively. A baseline block involving 6 trials (Pre) was completed prior to TMS measures. VT then involved 8 blocks of 8 trials (B1–B8). Participants completed each trial at their own pace, but they were instructed to focus on improving their speed and accuracy during each trial. A ‘skill’ score (see below) was calculated at the end of each block and displayed on a screen to provide feedback on performance.

### Data analysis

#### MEP data

MEP data were inspected visually and trials with muscle activity > 20 µV peak-to-peak amplitude in the 100 ms prior to TMS were rejected. MEP amplitude recorded in each trial was then quantified peak-to-peak and expressed in millivolts (mV). MEP amplitudes recorded during iTMS were averaged over 10 consecutive stimuli, resulting in a total of 18 blocks.

#### VT data

Skill scores were calculated for each block based on the movement speed and accuracy. Speed was measured by the average movement time (MT) for each trial. Accuracy was defined based on the error between the applied force and the force required to meet the center of the target. This was calculated for each of the 5 force peaks within a trial using the Euclidean distance, and then averaged over peaks to produce a trial error score. Skill scores were finally calculated using the following formula, as proposed by Reis *et al*. (2009).

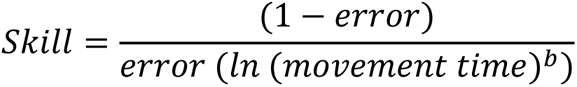

The dimensionless free b parameter has been shown to be insensitive to changes in performance, and thus was set at a consistent 1.627 (Stavrinos & Coxon, 2017).

#### EEG data

All preprocessing and subsequent analysis was performed according to previously reported procedures (Rogasch *et al*., 2017; Mutanen *et al*., 2020; Sasaki *et al*., 2022) using custom scripts on the MATLAB platform (R2019b, Mathworks, USA), in addition to EEGLAB (v2020.0) (Delorme & Makeig, 2004), TESA (v1.1.1.) (for review, see Rogasch *et al*., 2017) and Fieldtrip (v20200607) (Oostenveld *et al*., 2011) toolboxes. Data were epoched from −1500 ms to 2000 ms around the TMS trigger, baseline corrected from −500 ms to −5 ms and merged into a single file including both TMS (Pre, Post iTMS, and Post Train) and sham. Channels demonstrating persistent, large amplitude muscle activity or noise were manually removed, and the peak of the TMS artifact was removed by cutting the data from −2 to 10 ms and replacing it using cubic interpolation. The data was subsequently downsampled from 8 kHz to 500 Hz and epochs demonstrating bursts of muscle activity or electrode noise were manually removed. Interpolated data from −2 to 10 ms was then replaced with constant amplitude data (i.e., 0 s) and the conditions were split into two separate files (real TMS and sham). An initial independent component analysis (ICA) was run on each file using the FastICA algorithm (Hyvarinen & Oja, 2000), and a couple of independent components (IC’s) representing the tail of the TMS-evoked muscle artifact were removed (for review, see Rogasch *et al*., 2017). Constant amplitude data from −2 to 10 ms were then replaced with cubic interpolation prior to the application of band-pass (1–100 Hz) and notch (48–52 Hz) filtering (zero-phase 4^th^ order Butterworth filter implemented). In order to remove any additional decay artifacts still present after the first round of ICA, the source-estimate-utilizing noise-discarding (SOUND) algorithm was then applied; this approach estimates and removes artefactual components within source space, and also allows missing electrodes to be estimated and replaced (Mutanen *et al*., 2018). A regularization parameter of 0.1 was used and 5 iterations were completed. Following SOUND, data around the TMS pulse were again replaced with constant amplitude data prior to application of a second round of ICA, and IC’s associated with blinks, eye movements, electrode noise, and muscle activity were automatically identified using the TESA compselect function (default settings) and visually inspected prior to removal (for review, see Rogasch *et al*., 2017). Data around the TMS pulse were then replaced with cubic interpolation, and all channels were re-referenced to average prior to a final baseline correction (−500 ms to −5 ms).

### Statistical analysis

All analyses were performed using PASW statistics software version 28 (SPSS; IBM, Armonk, NY, USA) or Fieldtrip toolbox (EEG data only). All data were assessed using generalized linear mixed models (GLMM). Data distribution was initially assessed using Kolmogorov-Smirnov tests and Q-Q plots (Lo & Andrews, 2015; Puri & Hinder, 2022). These identified that VAS (all items) and iTMS intensity were normally distributed and could therefore be fit with a Gaussian distribution (i.e., linear mixed model). However, other TMS intensities, MEP amplitude, and VT data all showed negatively skewed distributions and were therefore modelled using a Gamma distribution with identity link function (Lo and Andrews, 2015). Each model involving MEP responses (raw MEP amplitude) used individual trial data, whereas all models included the maximal participant random effects structure. Model fit was assessed using the Akaike’s Information Criterion (AIC). Post hoc analysis of all significant main effects and interactions were performed using custom contrasts with Bonferroni correction, and significance was set at *P* < 0.05. All data are presented as estimated marginal means (EMM) and 95% confidence intervals (95% CI).

#### MEP data

One-factor GLMM analysis with repeated measures (GLMM_RM_) was used to compare baseline RMT, TS intensity, iTMS intensity, MEP_TS_, and MEP_iTMS_ between sessions (iTMS+VT, iTMS*_Sham_*+VT, and iTMS). For TS MEP amplitudes before and after interventions, two-factor GLMM_RM_ was used to compare values between sessions and time points (Pre, Post iTMS, and Post Train). Two-factor GLMM_RM_ was also used to compare MEP amplitudes during iTMS between sessions and blocks (B1–B18).

#### VT data

One-factor GLMM_RM_ was used to compare baseline error, MT, and skill between sessions (iTMS+VT, iTMS*_Sham_*+VT, and iTMS). Two-factor GLMM_RM_ was also used to compare error, MT, and skill between sessions (iTMS+VT and iTMS*_Sham_*+VT) and blocks (Pre, B1–B8).

#### TEP data

In an attempt to identify the elements of the EEG signal that were likely to be more contaminated by auditory and somatosensory inputs, the TEP produced by M1 stimulation was compared to the response generated by sham stimulation in both spatial (i.e., between electrodes at each time point) and temporal (i.e., across time points within each electrode) domains using the Spearman correlation coefficient (Biabani *et al*., 2019; Sasaki *et al*., 2021). Spatial analyses were conducted from −50 to 350 ms, whereas temporal analyses were averaged over early (15–60 ms), middle (60–180 ms) and late (180–280 ms) time periods (Sasaki *et al*., 2021). For both measures, correlation coefficients were converted to Z-values using Fisher’s transform prior to group analysis (Rocchi *et al*., 2020; Sasaki *et al*., 2021). Statistical significance was then determined using a one-sample permutation test (derived from 10,000 permutations) assessing the hypothesis that each Z-score was greater than zero (i.e., positive correlation), with the t_max_ method used to control the family-wise error rate for multiple comparisons (Fernandez *et al*., 2021). The Z-values were finally transformed back into their original form for display (Fernandez *et al*., 2021).

For data within each session, TEPs were compared between Pre and Post iTMS, Pre and Post Train, or Post iTMS and Post Train using cluster-based non-parametric permutation analysis. Furthermore, baseline TEPs were compared between sessions. Clusters were defined as two or more neighboring electrodes and 10,000 iterations were applied. A cluster was deemed significant if the cluster statistic exceeded *P* < 0.05 when compared with the permutation distribution. As correlation analysis demonstrated that TEPs were highly related to the response to sham stimulation from ∼60 ms post-stimulus (see Figure 7), comparisons between conditions were limited to the early TEP components, including N15 (10–15 ms), P30 (25–35 ms) and N45 (40–50 ms).

#### VAS data

Two-factor GLMM_RM_ was used to compare auditory intensity, scalp intensity, scalp area, and pain between sessions and stimulation types (TMS and ES).

## Results

All 16 participants completed the 3 sessions without any adverse events (mean time between sessions ± SD: S1–S2, 9.6 ± 3.7 days; S2–S3, 12.0 ± 8.0 days). A total of 2.8% and 5.8% of trials were removed from TS MEP and during iTMS MEP, respectively. Baseline characteristics for MEP and VT are compared between sessions in Table 1. Comparisons of RMT and TS intensity showed no differences between sessions (RMT: *F*_(2,45)_ = 2.747, *P* = 0.075; TS: *F*_(2,45)_ = 0.189, *P* = 0.828), but iTMS intensity was higher for iTMS*_Sham_*+VT than other sessions (*F*_(2,45)_ = 37.366, *P* < 0.001). Baseline TS and iTMS MEP amplitudes showed no differences between sessions (MEP_TS_: *F*_(2,690)_ = 0.362, *P* = 0.697; MEP_iTMS_: *F*_(2,477)_ = 1.593, *P* = 0.204). Furthermore, comparisons of baseline error, MT, and skill showed no differences between sessions (Error: *F*_(2,284)_ = 1.763, *P* = 0.173; MT: *F*_(2,281)_ = 0.010, *P* = 0.990; Skill: *F*_(2,281)_ = 1.751, *P* = 0.176).

**Table 1.** Baseline characteristics, corticospinal responses, and motor skills for each session.

|  | iTMS+VT | iTMS<sub>sham</sub>+VT | iTMS |
| --- | --- | --- | --- |
| RMT (%MSO) | 59.3 [54.3, 64.3] | 61.5 [56.5, 66.5] | 60.1 [55.1, 65.1] |
| TS (%MSO) | 71.9 [65.2, 78.6] | 72.7 [66.0, 79.4] | 71.8 [65.2, 78.5] |
| iTMS (%MSO) | 59.8 [54.9, 64.8]* | 71.2 [66.2, 76.1] | 61.2 [56.2, 66.1]* |
| MEP<sub>TS</sub> (mV) | 0.74 [0.61, 0.87] | 0.70 [0.57, 0.83] | 0.69 [0.56, 0.82] |
| MEP<sub>iTMS</sub> (mV) | 0.46 [0.31, 0.62] | 0.52 [0.36, 0.68] | 0.63 [0.47, 0.80] |
| Error (a.u.) | 0.18 [0.13, 0.22] | 0.21 [0.16, 0.26] | 0.22 [0.17, 0.27] |
| MT (sec) | 3.33 [2.83, 3.91] | 3.35 [2.85, 3.93] | 3.34 [2.84, 3.92] |
| Skill (a.u.) | 4.60 [3.38, 5.82] | 3.77 [2.59, 4.95] | 3.65 [2.46, 4.84] |

EMM [95% CI; lower, upper]. \*P < 0.05 compared to iTMS<sub>sham</sub>+VT. Abbreviations: iTMS<sub>sham</sub>, control I-wave periodicity repetitive transcranial magnetic stimulation; iTMS, I-wave periodicity repetitive transcranial magnetic stimulation; MEP, motor-evoked potential; MEP<sub>iTMS</sub>, MEP amplitude producing a response of ~0.5–1 mV by iTMS; MEP<sub>TS</sub>, MEP amplitude producing a response of ~0.5–1 mV by single pulse TMS; %MSO, %maximum stimulator output; MT, movement time; RMT, resting motor threshold; TS, test stimulus; VT, visuomotor task.

VAS for each item is compared between sessions and stimulus conditions in Table 2. Auditory intensity was not different between sessions (*F*_(1,90)_ = 2.301, *P* = 0.106), and there was no interaction between factors (*F*_(2,90)_ = 0.478, *P* = 0.621). However, values were higher for TMS than ES (*F*_(1,90)_ = 7.723, *P* = 0.007). Furthermore, scalp area was not different between sessions (*F*_(1,88)_ = 2.346, *P* = 0.709), and there was no interaction between factors (*F*_(2,88)_ = 0.623, *P* = 0.539). However, values were higher for TMS than ES (*F*_(1,88)_ = 8.500, *P* = 0.005). No differences were found for other items (*P* > 0.32).

**Table. 2.** VAS between TMS and ES.

|  |  | iTMS+VT | iTMS<sub>sham</sub>+VT | iTMS |
| --- | --- | --- | --- | --- |
| Auditory intensity | TMS | 3.0 [2.2, 3.8] | 2.6 [1.7, 3.4] | 3.8 [2.9, 4.6] |
|  | ES | 2.3 [1.4, 3.1]* | 2.1 [1.2, 2.9]* | 2.9 [2.0, 3.7]* |
| Scalp intensity | TMS | 2.8 [1.6, 4.0] | 2.6 [1.4, 3.8] | 3.6 [2.4, 4.7] |
|  | ES | 2.7 [1.6, 3.9] | 2.9 [1.7, 4.1] | 3.1 [1.9, 4.2] |
| Stimulation area | TMS | 2.9 [1.9, 3.9] | 3.3 [2.3, 4.3] | 3.1 [2.1, 4.1] |
|  | ES | 1.9 [0.9, 2.9]* | 2.1 [1.1, 3.1]* | 1.7 [0.7, 2.7]* |
| Pain | TMS | 0.9 [0.3, 1.6] | 0.6 [-0.1, 1.2] | 0.6 [-0.1, 1.2] |
|  | ES | 0.9 [0.2, 1.5] | 0.9 [0.2, 1.5] | 0.6 [-0.1, 1.2] |

EMM [95% CI; lower, upper]. \*P < 0.05 compared to TMS. Abbreviations: iTMS<sub>sham</sub>, control I-wave periodicity repetitive transcranial magnetic stimulation; ES, electrical stimulation; iTMS, I-wave periodicity repetitive transcranial magnetic stimulation; VT, visuomotor task.

### Effects of iTMS on corticospinal excitability

Figure 2A shows changes in MEP amplitude during iTMS. No difference was found between sessions (*F*_(2,8085)_ = 1.054, *P* = 0.349), and there was no interaction between factors (*F*_(34,8085)_ = 0.739, *P* = 0.865). However, values varied over blocks (*F*_(17,8085)_ = 1.881, *P* = 0.015), with post-hoc comparisons showing increased amplitude during block 17 relative to block 1 (*P* = 0.049). TS MEP amplitudes before and after iTMS and VT are shown in Figure 2B. MEP amplitudes were not different between sessions (*F*_(2,2090)_ = 0.554, *P* = 0.575) or time points (*F*_(2,2090)_ = 1.557, *P* = 0.211) and there was no interaction between factors (*F*_(4,2090)_ = 1.251, *P* = 0.287).

**Figure 2.**
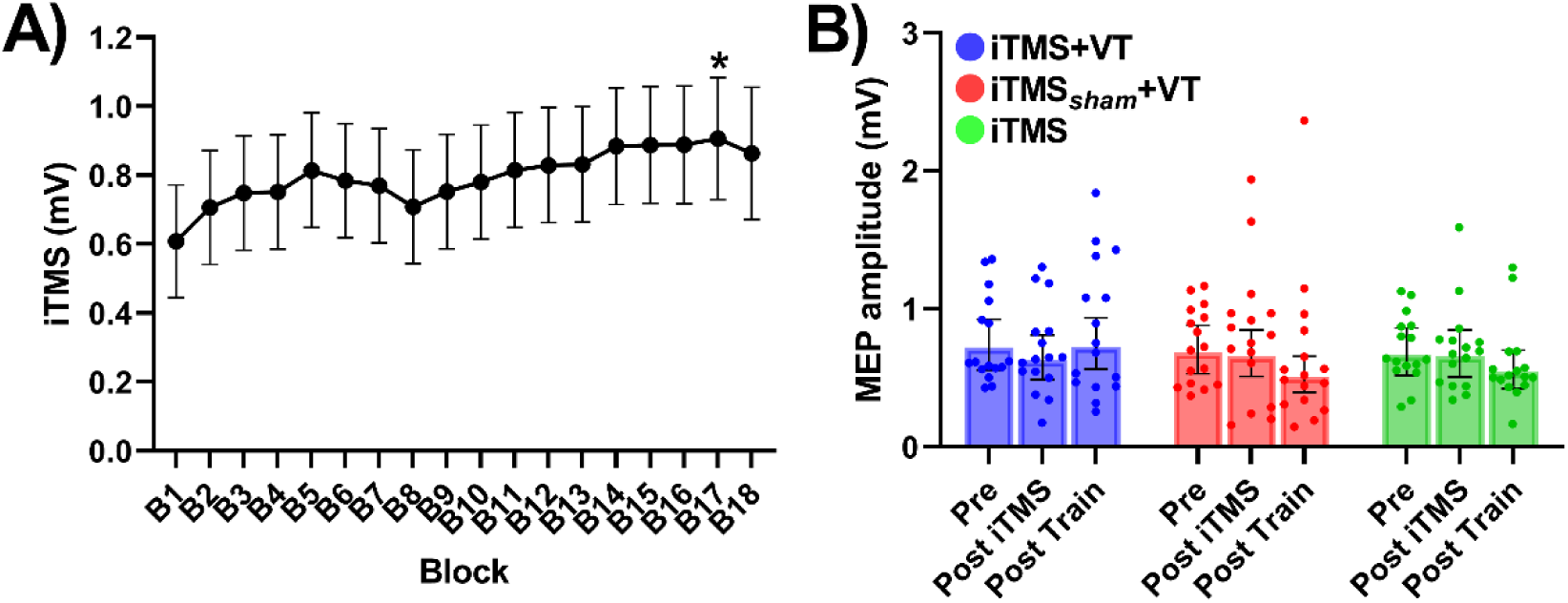
Corticospinal excitability changes by iTMS and VT. (A) MEP amplitudes during iTMS, averaged over 10 consecutive MEP trials. (B) TS MEP amplitudes before and after iTMS and VT. *P < 0.05 compared to B1. EMM ± 95% CI. Abbreviations; B, block; iTMS_sham_, control I-wave periodicity repetitive transcranial magnetic stimulation; iTMS, I-wave periodicity repetitive transcranial magnetic stimulation; MEP, motor-evoked potential; VT, visuomotor task.

### Effects of iTMS on visuomotor training

Performance during VT is shown in figure 3. Error was not different between sessions (*F*_(1,2219)_ = 1.923, *P* = 0.166), and there was no interaction between factors (*F*_(8,2219)_ = 0.343, *P* = 0.949). However, error varied over blocks (*F*_(8,2219)_ = 3.613, *P* < 0.001), with *post-hoc* comparisons showing decreased error during training (i.e., block 1–8) relative to baseline (all *P* < 0.02)(Fig 3A). MT was not different between sessions (*F*_(1,2198)_ = 0.828, *P* = 0.363) and there was no interaction between factors (*F*_(8,2198)_ = 0.768, *P* = 0.631). However, MT varied over blocks (*F*_(8,2298)_ = 19.806, *P* < 0.001), with *post-hoc* comparisons showing decreased MT during block 2–8 relative to Pre (all *P* < 0.001)(Fig 3B). Furthermore, skill varied between sessions (*F*_(1,2205)_ = 6.044, *P* = 0.014), with *post-hoc* comparisons showing greater skill for iTMS+VT relative to iTMS*_Sham_*+VT (*P* = 0.014)(Fig 3C). Skill also varied over blocks (*F*_(8,2205)_ = 26.844, *P* < 0.001), with *post-hoc* comparisons showing increased skill during block 1–8 relative to Pre (all *P* < 0.002). However, there was no interaction between factors (*F*_(8,2205)_ = 0.390, *P* = 0.926).

**Figure 3.**
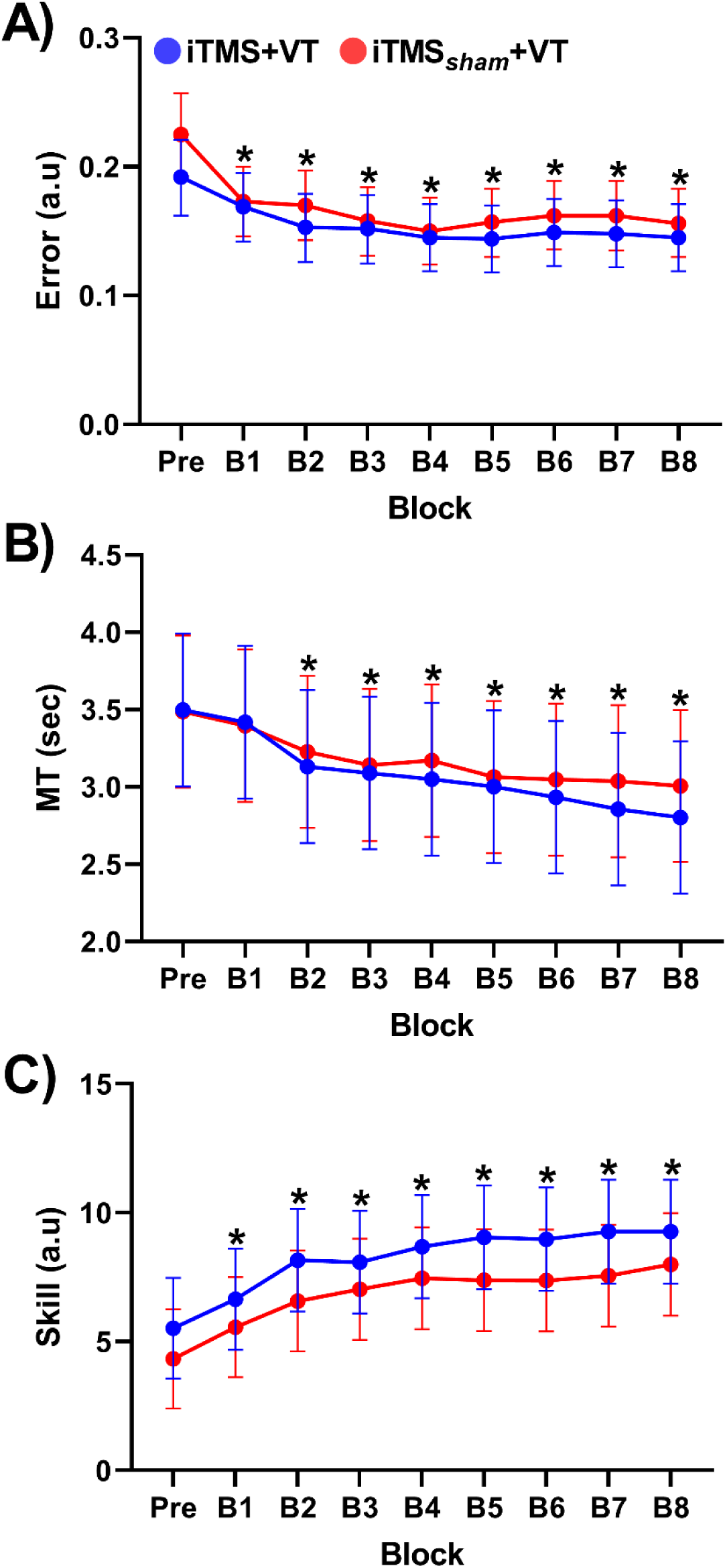
Changes in motor skills over blocks. Panels (A, B, C) represent error, MT, and skill before and after iTMS, respectively. *P < 0.05 compared to Pre. EMM ± 95% CI. Abbreviations; B, block; iTMS_sham_, control I-wave periodicity repetitive paired-pulse transcranial magnetic stimulation; iTMS, repetitive I-wave periodicity paired-pulse transcranial magnetic stimulation; MT, movement time; VT, visuomotor task.

Given the differences in skill across blocks that included the baseline timepoint, the analysis of motor performance measures was repeated using data that were expressed as a percentage of baseline. Using this approach, Error was no longer different between blocks (*F*_(7,2029)_ = 0.784, *P* = 0.600), whereas skill was no longer different between sessions (*F*_(1,2016)_ = 2.587, *P* = 0.108). All other results were consistent with the original analysis of non-normalised data.

### TEP preprocessing and correlation analysis

The average number of channels, epochs and IC’s removed during each step of the preprocessing pipeline are shown in Table 3. Figures 4, 5 and 6 show grand-average TEP waveforms elicited by M1 and electrical stimulation, whereas Figure 7 shows correlation coefficients resulting from comparisons between M1 and electrical stimulation in both spatial (Figure 7A, B, C) and temporal (Figure 7D) domains. For all sessions, spatial correlations mainly identified significant relationships between conditions at the Late period. In support of this, results of the temporal correlations suggested that the two signals were largely unrelated within the Early period, but became highly correlated across the scalp in the Mid and Late periods. These results suggest that, although the early TEP response was likely to be less contaminated by sensory inputs, signal within the Mid and Late periods were likely to be heavily contaminated. Consequently, all statistical analyses of TEP amplitude were limited to the early period.

**Figure 4.**
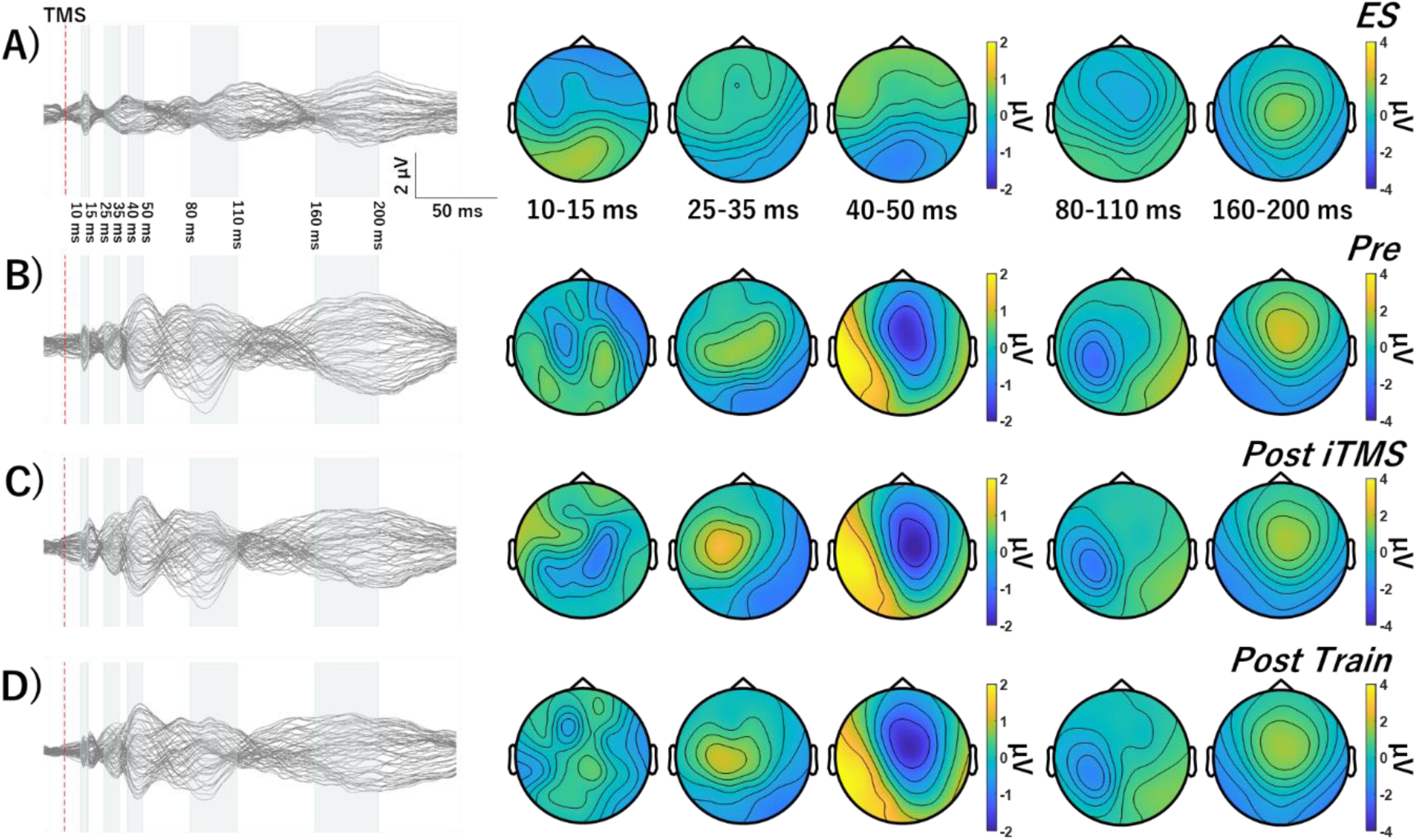
Grand average TEP waveforms and topographies in iTMS+VT session. (A, B, C) ES before iTMS (A) and M1 stimulation before and after iTMS and VT (B, C, D). Baseline TEP waveforms show several typical TEP components, named as N15, P30, P45, N100, and P180. Abbreviations; ES, electrical stimulation; TMS, transcranial magnetic stimulation; iTMS, I-wave periodicity repetitive paired-pulse transcranial magnetic stimulation.

**Figure 5.**
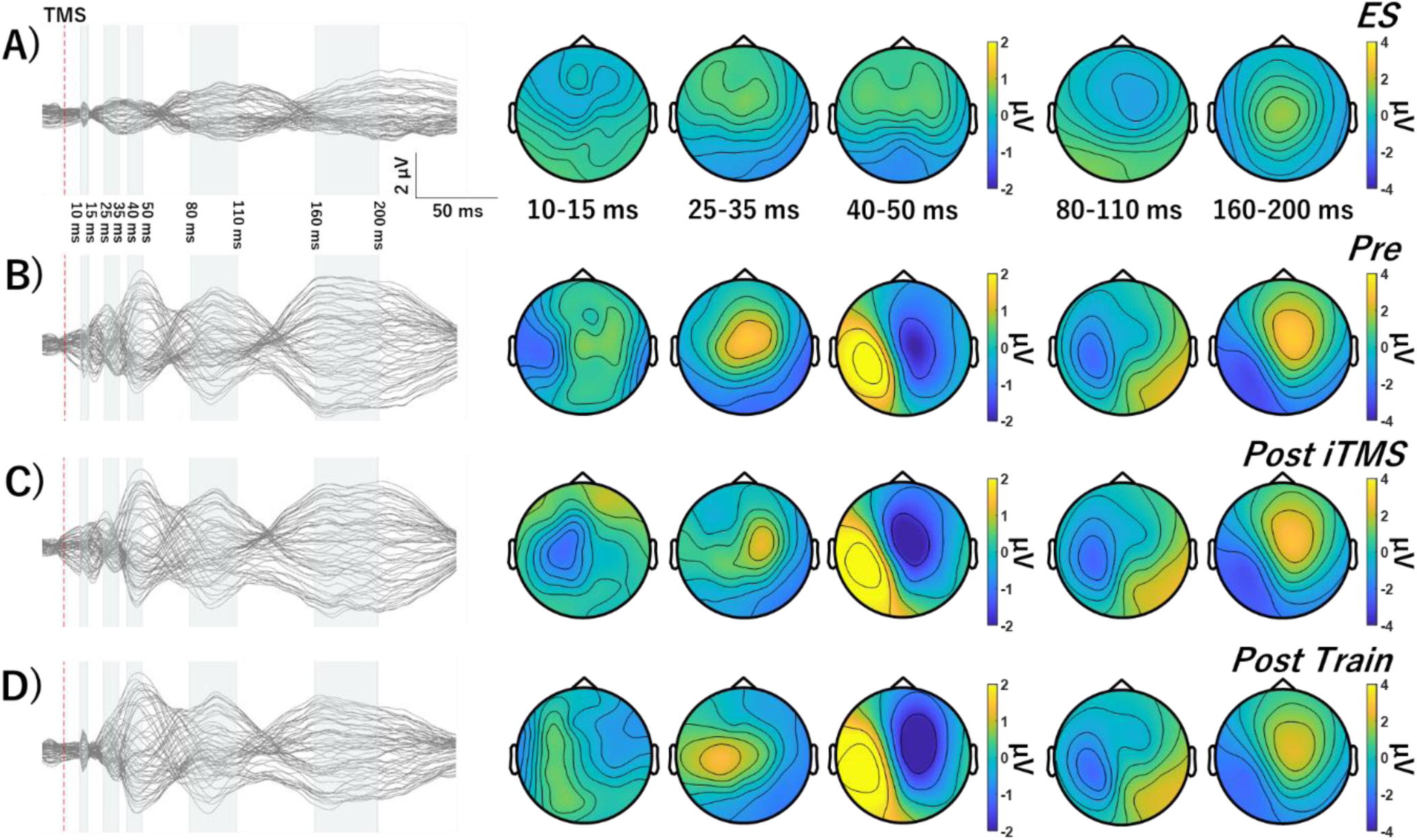
Grand average TEP waveforms and topographies in iTMS_Sham_+VT session. (A, B, C) ES before iTMS (A) and M1 stimulation before and after iTMS_sham_ and VT (B, C, D). Baseline TEP waveforms show several typical TEP components, named as N15, P30, P45, N100, and P180. Abbreviations; iTMS_sham_, control I-wave periodicity repetitive paired-pulse transcranial magnetic stimulation; ES, electrical stimulation; TMS, transcranial magnetic stimulation.

**Figure 6.**
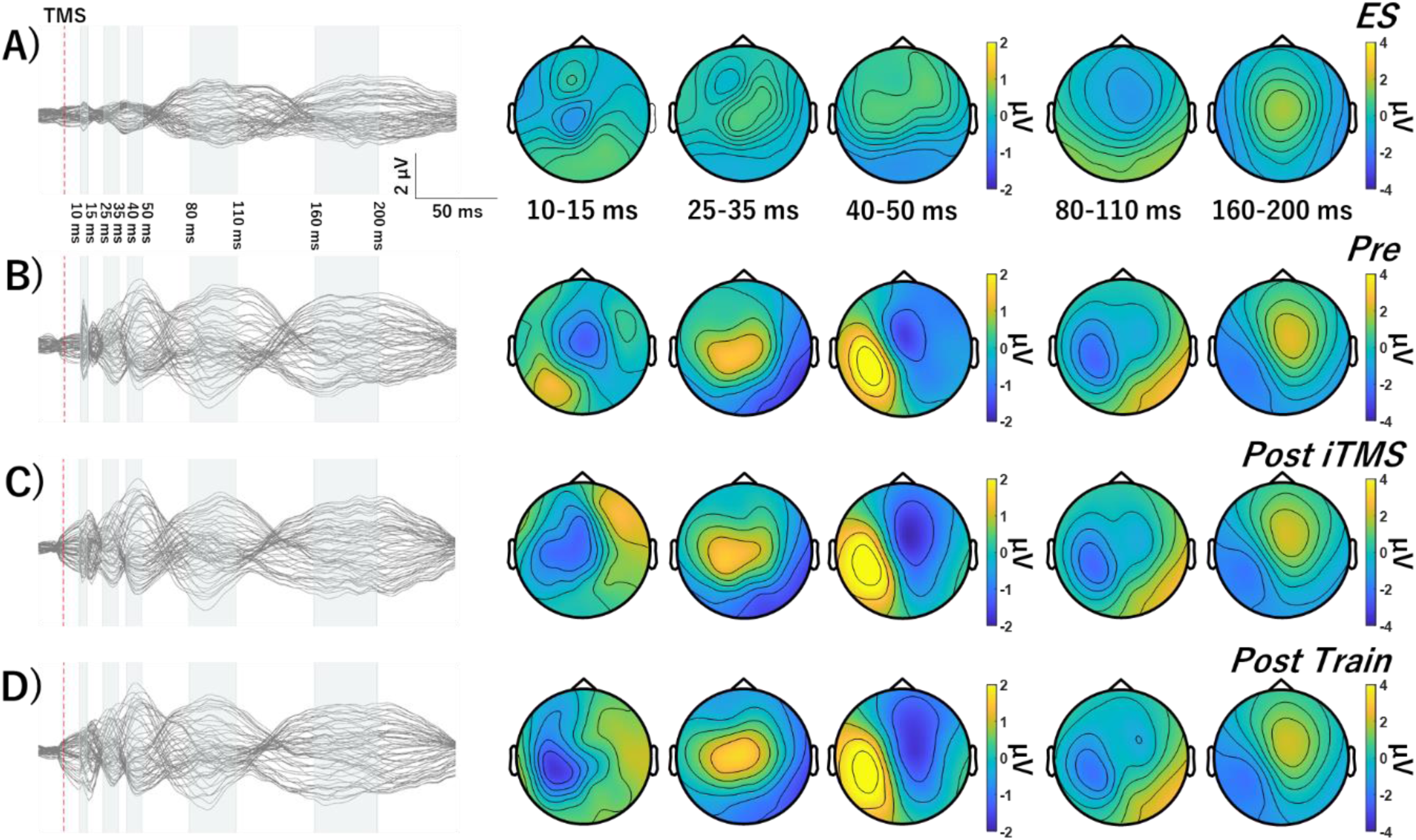
Grand average TEP waveforms and topographies in iTMS session. (A, B, C) ES before iTMS (A) and M1 stimulation before and after iTMS and 15 min break (B, C, D). Baseline TEP waveforms show several typical TEP components, named as N15, P30, P45, N100, and P180. Abbreviations; ES, electrical stimulation; TMS, transcranial magnetic stimulation; iTMS, I-wave periodicity repetitive paired-pulse transcranial magnetic stimulation.

**Figure 7.**
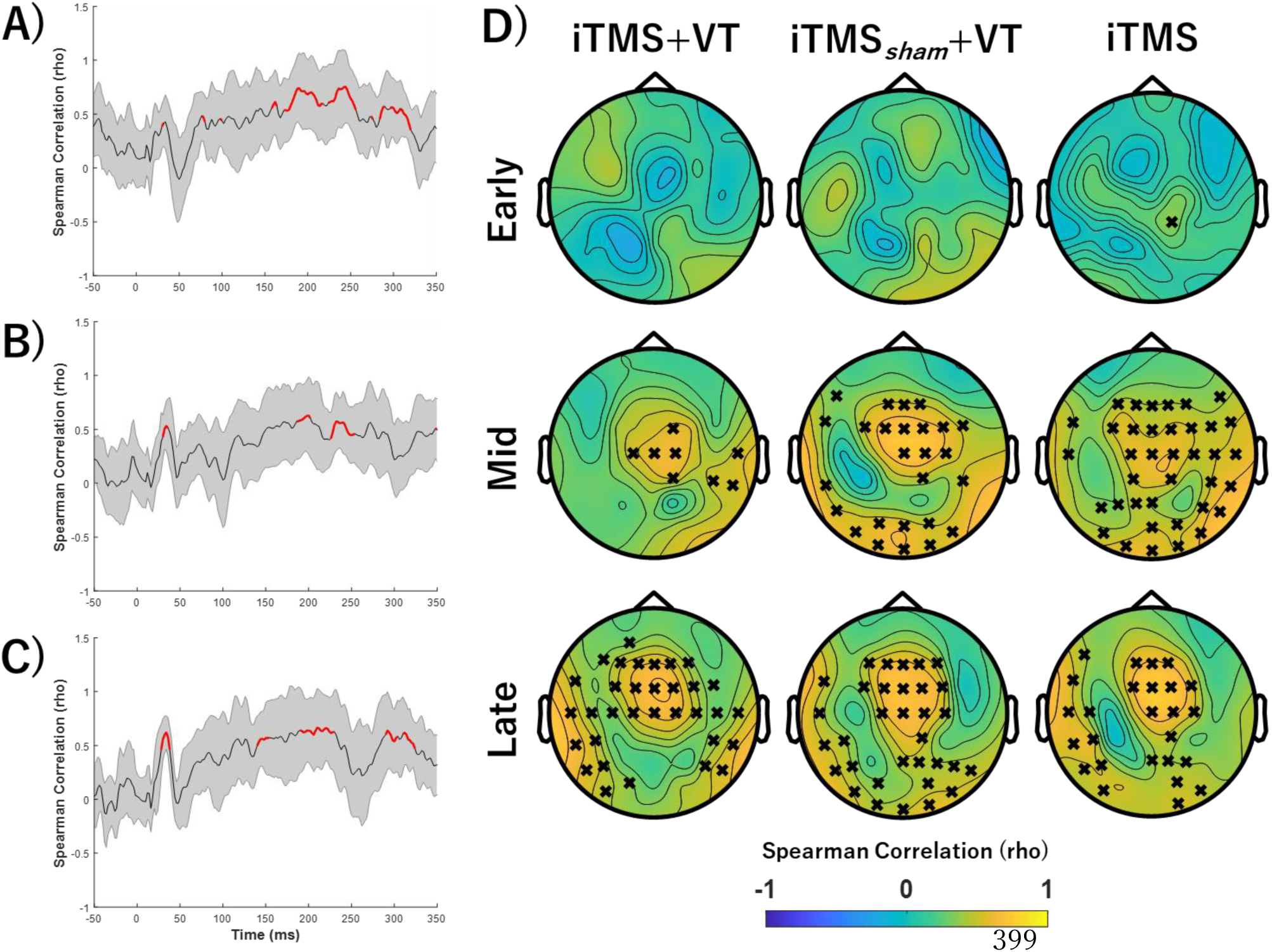
TEPs and sensory correlations. (A, B, C) Spatial correlations between EEG response to M1 and electrical stimulation in iTMS+VT (A) and iTMS_sham_+VT (B), and iTMS (C) sessions across all EEG electrodes. Red line segments indicate time periods that are significantly related between stimulation conditions. (D) Temporal correlations between EEG response to M1 and electrical stimulation during Early (15–60 ms), Mid (60–180 ms) and Late (180–280 ms) time periods. Black crosses show that electrodes were significantly correlated between conditions. Abbreviations; iTMS_sham_, control I-wave periodicity repetitive paired-pulse transcranial magnetic stimulation; iTMS, I-wave periodicity repetitive paired-pulse transcranial magnetic stimulation.

**Table 3.** Number of channels, epochs, and independent components removed during.

|  | iTMS+VT | iTMS<sub>sham</sub>+VT | iTMS |
| --- | --- | --- | --- |
| Channels (TS_pre) | $0.38 \pm 0.17$ | $0.38 \pm 0.21$ | $0.44 \pm 0.23$ |
| Channels (TS_Post iTMS) | $0.38 \pm 0.17$ | $0.38 \pm 0.21$ | $0.44 \pm 0.23$ |
| Channels (TS_Post Train) | $0.38 \pm 0.17$ | $0.38 \pm 0.21$ | $0.44 \pm 0.23$ |
| Channels (sham) | $0.38 \pm 0.17$ | $0.38 \pm 0.21$ | $0.31 \pm 0.21$ |
| Epoch (TS_pre) | $1.94 \pm 0.42$ | $2.13 \pm 0.48$ | $1.75 \pm 0.55$ |
| Epoch (TS_Post iTMS) | $4.06 \pm 1.14$ | $2.25 \pm 0.48$ | $2.00 \pm 0.49$ |
| Epoch (TS_Post Train) | $1.88 \pm 0.75$ | $2.81 \pm 1.09$ | $2.13 \pm 0.87$ |
| Epoch (sham) | $1.81 \pm 0.38$ | $2.44 \pm 0.69$ | $2.94 \pm 0.80$ |
| ICA1 (TS) | $2.75 \pm 0.40$ | $2.56 \pm 0.44$ | $2.75 \pm 0.43$ |
| ICA1 (sham) | $1.56 \pm 0.15$ | $1.56 \pm 0.15$ | $1.50 \pm 0.15$ |
| ICA2 (TS) | $7.25 \pm 0.79$ | $7.63 \pm 0.81$ | $7.13 \pm 0.82$ |
| ICA2 (sham) | $5.50 \pm 0.59$ | $4.63 \pm 0.47$ | $5.56 \pm 0.54$ |

Mean ± SEM. Abbreviations; iTMS<sub>sham</sub>, control I-wave periodicity repetitive paired-pulse transcranial magnetic stimulation; iTMS, I-wave periodicity repetitive paired-pulse transcranial magnetic stimulation; ICA, independent component analysis; TS, test stimulus; VT, visuomotor task.

### Changes in cortical excitability before and after interventions

Baseline TEP components were not different between sessions (all *P* > 0.08). For each iTMS+VT and iTMS session, there were no differences between time points (all *P* > 0.15). For iTMS*_Sham_*+VT, comparisons of P30 between Pre and Post Train identified significant negative and (*P* = 0.028) and positive clusters *(P* = 0.042). Comparisons of P30 between Post iTMS and Post Train also identified a significant positive cluster (*P* = 0.033) (Figure 8).

**Figure 8.**
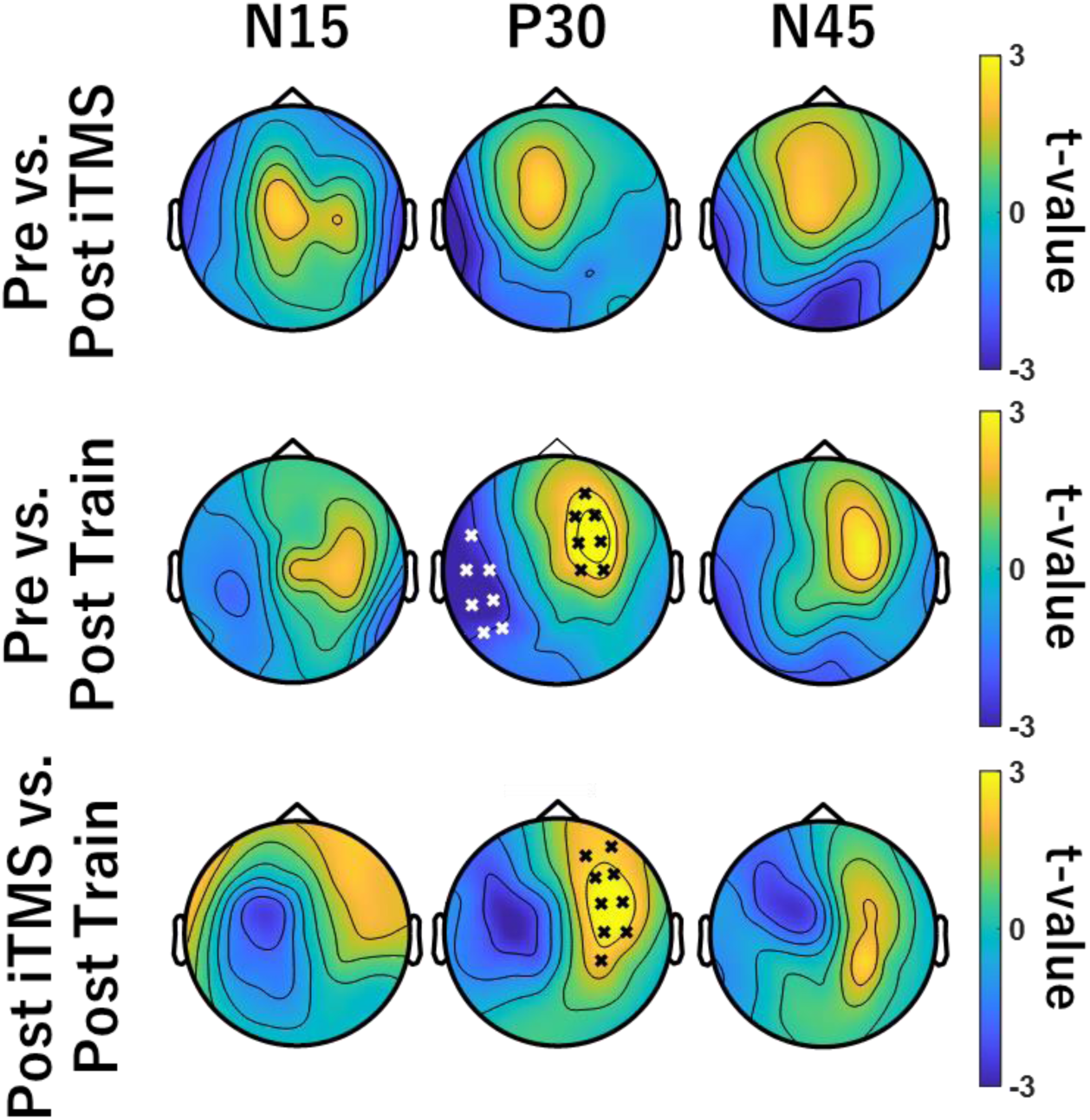
Comparison of TEPs between Pre and Post in iTMS_Sham_+VT session. These topographies represent cluster-based permutation t-test comparing the TEPs amplitudes before and after iTMS_sham_ immediately (top row), before iTMS_sham_ and after VT (middle row), after iTMS_sham_ immediately and after VT (bottom row). Black and white crosses show significant clusters between Pre- and Post Train- or Post iTMS- and Post Train-P30 amplitude.

However, no differences were found for the N15 and N45 (all *P* = 1).

## Discussion

Within the current study, we aimed to further characterise the neurophysiological processes that underpin beneficial effects of iTMS on motor learning. To achieve this, TEPs were recorded before and after a visuomotor adaptation task that was practiced in isolation, or following application of iTMS. While skill increased in response to training, the magnitude of this effect was not different between priming conditions, suggesting that iTMS was ineffective as a priming intervention. However, iTMS also failed to induce the expected potentiation of MEP amplitude, complicating interpretation of the response to training.

Despite this, differential effects on TEP amplitude suggested that training produced changes in cortical activity that were cancelled by priming.

### Skill acquisition, corticospinal excitability and priming

Previous work has reported that, when applied prior to training, a neuromodulatory NIBS intervention can improve acquisition of a novel motor skill (e.g., Jung & Ziemann, 2009). Within this construct, NIBS-dependent modulation of motor network activity is thought to generate a neural environment that is more amenable to the neuroplastic changes required to learn new patterns of motor behaviour (Müller-Dahlhaus & Ziemann, 2015). This has been supported by studies showing that priming-dependent modulation of motor learning is accompanied by related changes in motor cortical excitability (Ziemann *et al*., 2004; Jung & Ziemann, 2009). Within the current study, visuomotor training resulted in improved skill levels that are consistent with previous work from our group (Opie *et al*., 2020; Hand *et al*., 2021; Hand *et al*., 2023) and others (Reis *et al*., 2009; Ho *et al*., 2022). While skill was significantly greater in the iTMS+VT condition, examination of normalised data showed this stemmed from baseline differences in performance (see *Results* and Fig 3C). In addition, MEP measures of corticospinal excitability were also unchanged by priming or training.

Taken together, our results therefore suggest that iTMS in the current study was unable to influence skill acquisition or corticospinal excitability (when assessed with MEPs). While overt changes in excitability are not a prerequisite for induction of metaplastic effects (e.g., (Ni *et al*., 2014; Fujiyama *et al*., 2017), the lack of change in corticospinal excitability nonetheless makes it difficult to interpret the training results. In particular, our recent work showed that iTMS increased MEP amplitude and improved SVIPT acquisition in both young and older adults (Hand *et al*., 2023), demonstrating the utility of this approach. This variability demonstrates that further examination of the factors driving functionally relevant effects of iTMS is required.

Given the similarity of the methodology between our current and previous (Hand *et al*., 2023) findings (including the same research environment and protocols), the extent of the divergence in results is surprising. A minor discrepancy between the studies was that the iTMS ISI differed by 0.1 ms, possibly contributing to variability. However, it can be expected that the timing of I-waves within individual participants varied by more than 0.1 ms (Sewerin *et al*., 2011). Consequently, it seems that the minor difference in ISI between studies would explain less variance than can be accounted for by the fixed ISI (relative to I-wave timing within individuals), and certainly wouldn’t account for the divergent findings of these studies. A more likely explanation is that the results reported here further demonstrate the variability that is being increasingly recognised within the field, particularly with respect to replication of canonical effects. For example, there is a growing literature that reports negative findings with respect to the effects of both neuromodulatory interventions (Hamada *et al*., 2013; López-Alonso *et al*., 2014; Wiethoff *et al*., 2014; Jonker *et al*., 2021) and motor training (Bestmann & Krakauer, 2015) on MEP amplitude, in addition to the effect of priming stimulation on motor learning (Lopez-Alonso *et al*., 2018; Sasaki *et al*., 2018).

The factors driving this variability are likely to be multifactorial; these have been covered in detail elsewhere (Ridding & Ziemann, 2010), but are known to include attention, cortisol levels (Sale *et al*., 2007; Sale *et al*., 2008), genetics (Cheeran *et al*., 2008), physical activity (Cirillo *et al*., 2009), chronotype (Salehinejad *et al*., 2021) and neural activity (Zrenner *et al*., 2022), in addition to the many potential methodological sources of variability (including statistical)(for review, see Guerra *et al*., 2020). An additional point that the current study can speak to (to some extent) is the way in which outcomes are assessed. For example, while MEPs were insensitive to the intervention applied here, TEPs were instead altered by training (see below). We do not mean to suggest that TEPs should be considered a superior approach; indeed, these responses are still heavily encumbered by methodological limitations, and their interpretation is being actively developed. Nonetheless, the contrast between findings reported here demonstrates the potential for alternative outcome measures to influence our results.

Control iTMS within the current study involved single-pulse stimulation applied with the same frequency and duration as real iTMS. This approach has been used by previous iTMS studies, which reported no change in MEPs during or after application (Silbert *et al*., 2011; Teo *et al*., 2012). In contrast to this, we found an apparent increase in MEP amplitude during application of control iTMS (data not shown). Although inconsistent with previous iTMS studies, other work has shown that there can be cumulative effects of single-pulse TMS over a period comparable to the application of iTMS (Pellicciari *et al*., 2016). While the specific reason this was apparent in the current but not previous studies remains unclear, it nonetheless demonstrates the need for an improved sham paradigm for iTMS. We have previously used sham stimulation that involved paired-pulse stimuli with ISIs associated with non-facilitatory periods of the I-wave recruitment profile, the order of which are pseudorandomised between trials (Liao *et al*., 2022). While this appears to be a promising approach, it has only been applied during application of cerebellar tDCS and will therefore need to be verified during isolated application to M1.

### Effects of motor training on cortical reactivity are removed following iTMS

Consistent with previous work (Biabani *et al*., 2019; Sasaki *et al*., 2022), correlations between real and sham TEPs suggested sensory contamination of late TEP components (Fig 7), and statistical comparisons between conditions were therefore restricted to early peaks thought to be less influenced by sensory input (i.e., N15, P30, and P45)(Conde *et al*., 2019; Gordon *et al*., 2021). While N15 and N45 were unchanged in any condition, P30 was found to vary in response to motor training alone (i.e., iTMS*_Sham_* + VT session). Specifically, amplitude was increased and more lateralized over ipsilateral central electrodes (Figs 5 & 8). While previous work has used TMS-EEG to investigate changes in cortical reactivity associated with visuomotor adaptation (Koch *et al*., 2020; Taga *et al*., 2021), effects of learning were limited to the later peaks that are associated with increased contamination from sensory input (Biabani *et al*., 2019). Consequently, as far as we are aware, the current study is the first to report a modulation of the early TEP peaks following visuomotor training. The P30 has been associated with local excitatory and inhibitory processes (Cash *et al*., 2017; Sasaki *et al*., 2021), and its modulation during training is therefore consistent with neural changes driven by motor learning (for review, see Dayan & Cohen, 2011). Interestingly, these changes were apparent despite MEPs being unaffected by learning, suggesting that TEP-based measures of cortical reactivity may be a more sensitive index of the neurophysiological response to training. However, it will be important for future work to investigate the test-retest reliability of this outcome to demonstrate its relevance to motor learning.

Whereas training alone resulted in a modulation of the TEP, this effect was removed when training was primed by iTMS. One explanation for this could be that priming stimulation interfered with the neuronal processes recruited by training. We recently reported effects of iTMS on TEPs that would generally be considered as beneficial to the neurophysiological processes associated with learning (i.e., disinhibition of local intracortical circuits; Ziemann *et al*., 2001; Sasaki *et al*., 2023) and it is therefore unclear why this would be the case.

However, the timing of this disinhibition is likely to be important (Ziemann & Siebner, 2008), and its application prior to learning may have resulted in metaplastic effects that interfered with the brains response to training. Nonetheless, these neurophysiological effects failed to influence the functional response to training. A question that stems from this is whether the cortical effects of priming: *(1)* failed to exceed some threshold required to influence learning or *(2)* were not directly relevant to learning/ were not conducive to improving learning. While the former option would suggest that increasing the strength of the priming stimulus (e.g., higher intensities, longer duration, paired priming blocks) may facilitate an impact on learning, the latter may instead imply that different priming would be needed, perhaps targeting other nodes of the motor network. The current study is unable to differentiate between these options and it will be important for future research to investigate them further.

In conclusion, the current study aimed to further investigate the neurophysiological effects of iTMS on cortical excitability and motor learning. Against expectations, the normally robust effects of iTMS on MEP amplitude were absent, training failed to modulate corticospinal excitability, and priming did not influence motor learning. In contrast, the P30 was modulated by motor learning, and this effect was removed when training was preceded by priming iTMS. While this suggests that priming was able to influence the cortical response to training, it remains unclear why this failed to impact learning.

## Abbreviations

EEG: electroencephalography
EMG: Surface electromyography
EMM: estimated marginal means
ES: Electrical stimulation
FDI: First dorsal interosseous
GLMM: generalised linear mixed models
ICA: independent component analysis
iTMS: Repetitive paired-pulse transcranial magnetic stimulation
M1: primary motor cortex
MEP: motor-evoked potentials
MSO: Maximum stimulator output
MT: movement time
MVC: Maximal voluntary contraction
PAS: Paired-associative stimulation
RMT: Resting motor threshold
TEP: TMS-evoked potential
TMS: Transcranial magnetic stimulation
TS: test stimulus
VAS: visual analog scale
VT: Visuomotor task

